# Perirhinal input to neocortical layer 1 controls learning

**DOI:** 10.1101/713883

**Authors:** Guy Doron, Jiyun N. Shin, Naoya Takahashi, Christina Bocklisch, Salina Skenderi, Moritz Drüke, Lisa de Mont, Maria Toumazo, Moritz von Heimendahl, Michael Brecht, Richard Naud, Matthew E. Larkum

**Affiliations:** Institute for Biology, Humboldt University of Berlin, D-10117 Berlin, Germany; Bernstein Center for Computational Neuroscience, Humboldt University of Berlin, D-10115 Berlin, Germany; uOttawa Brain and Mind Institute, Department of Cellular and Molecular Medicine, University of Ottawa, Ottawa, Ontario, K1H 8M5, Canada

## Abstract

Signals sent back to the neocortex from the hippocampus control the long-term storage of memories in the neocortex^1,2^, but the cellular mechanisms underlying this process remain elusive. Here, we show that learning is controlled by specific medial-temporal input to neocortical layer 1. To show this we used direct cortical microstimulation detection task that allowed the precise region of learning to be examined and manipulated. Chemogenetically suppressing the last stage of the medial temporal loop, i.e. perirhinal cortex input to neocortical layer 1, profoundly disrupted early memory formation but had no effect on behavior in trained animals. The learning involved the emergence of a small population of layer 5 pyramidal neurons (~10%) with significantly increased firing involving high-frequency bursts of action potentials that were also blocked by suppression of perirhinal input. Moreover, we found that dendritic excitability was correspondingly enhanced in a similarly-sized population of pyramidal neurons and suppression of dendritic activity via optogenetic activation of dendrite-targeting inhibitory neurons also suppressed learning. Finally, single-cell stimulation of cortical layer 5 pyramidal neurons showed that burst but not regular firing retrieved previously learned behavior. We conclude that the medial temporal input to the neocortex controls learning through a process in L1 that elevates dendritic calcium and promotes burst firing.

## Results

The distributed nature of long-term memory formation in the cortex has challenged research into the underlying mechanisms. For hippocampal-independent learning paradigms there is converging evidence to suggest that cortical layer 1 (L1) is a locus for plasticity^3–6^ involving activity in the distal tuft dendrites of pyramidal neurons that innervate L1^4,7–9^. Far less is known about the mechanisms underlying hippocampal-dependent memory formation in the cortex. Application of the retrograde tracer, Fast Blue, to L1 of primary somatosensory cortex (S1), revealed labeled cells in the deep layers of the perirhinal cortex (Fig. 1a, bottom left). Conversely, expression of ChR2-EYFP via a viral vector (AAV) injected into the deep layers of the perirhinal cortex densely labeled axons in L1 of S1 (Fig. 1a, bottom right), confirming that the perirhinal cortex is the last station in the medial temporal loop before the primary somatosensory neocortex in rodents (Fig. 1a, right and Extended Fig. 1)^2,10,11^.

**Figure 1.**
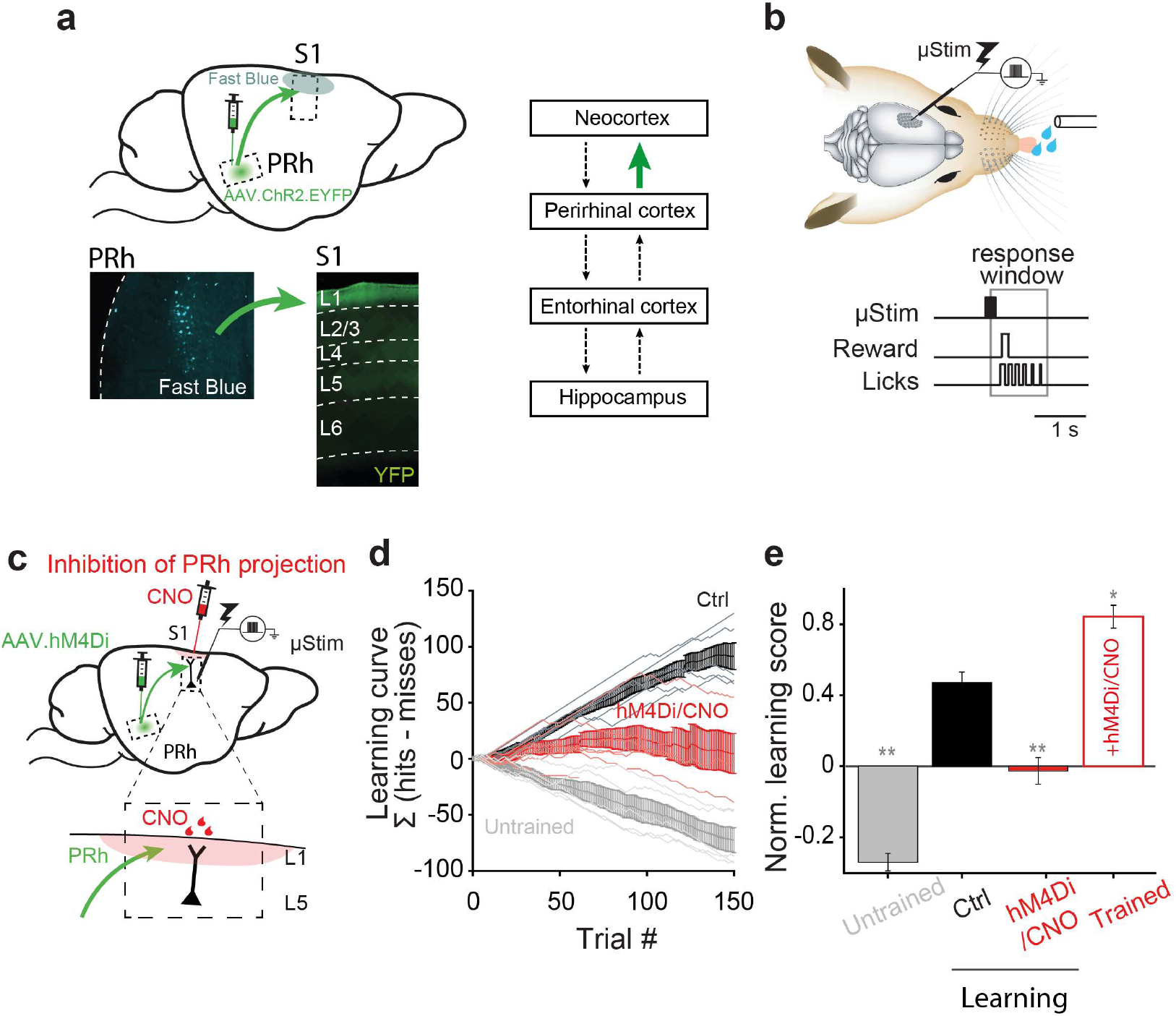
Perirhinal projection to neocortical L1 is necessary for learning a μStim task. **a**, Left, sagittal view of the rodent brain showing perirhinal projection to primary somatosensory cortex (S1). Lower left, retrograde tracing. Deep layer neurons in perirhinal cortex (PRh) were labeled with Fast Blue after application of dye to L1 of S1. Lower right, anterograde tracing. ChR2/EYFP labeled axons of PRh project strongly to L1 of S1. Right, simplified connectivity map between the neocortex, perirhinal cortex, entorhinal cortex and hippocampus. Investigated connection is highlighted in green. Note that not all the connections are shown here. **b**, Schematic of μStim detection task. Tungsten electrode was placed in L5 of S1. Animals learned to respond by licking within a 1.1-s window following μStim. **c**, Schematic of chemogenetic silencing of PRh axons in L1 of S1 during μStim task. AAV.hM4Di was injected to PRh and CNO was applied in superficial layer of S1 before μStim task (see Methods). Inset, enlarged view of superficial layer of S1. Red shade represents CNO effective area. **d**, Cumulative learning curve of control mice (black), mice with PRh axonal suppression (hM4Di; red) and untrained mice (grey) during 150 trials in the first session. Light lines represent individual mouse. Bold lines with error bars represent the mean and SEM, respectively, of each group. **e**, Last value of cumulative learning curve normalized by total number of trials (Norm. learning score) during learning and after learning (Trained). Wilcoxon rank-sum test against control, *p<0.05, **p<0.01.

In order to examine the influence of perirhinal cortex on memory formation in neocortex, we adapted a fast-learning, associative and cortex-dependent task^12^. Rodents were trained to report short (200 ms) trains of direct electrical microstimulation (μStim) pulses in layer 5 (L5) of S1 (Fig. 1b) where μStim detection threshold is lowest^12^. Animals initially received a block (5 repetitions) of μStim paired with the reward (sweetened water) regardless of their licking responses. Following a brief pairing period (1-2 blocks), the reward became available if the animal actively licked within a response window of 100–1200 ms following μStim onset (Fig. 1b). Animals learned this task extremely quickly during the first training session and became experts after about 3 training sessions (see Methods). Ipsilateral injections of lidocaine in the hippocampus showed that this task is hippocampus-dependent (Extended Fig. 2). Making behavior contingent on μStim of S1 allowed us to precisely define the area of interest and the temporal window in order to examine the underlying neuronal mechanisms of memory formation. Moreover, it allowed us to precisely target the perirhinal projection to L1 of S1. We chose a chemogenetic approach to down-regulate synaptic transmission^13^ at the axon terminals of perirhinal long-range projecting neurons without influencing the hippocampus and parahippocampal regions. Here, we expressed hM4Di receptors (inhibitory designer receptors exclusively activated by a designer drug, DREADD^14^) in the perirhinal cortex of mice (Fig. 1c). The axon terminals in S1 were inhibited by application of clozapine-N-oxide (CNO, 10 μM), injected in to L1 above the stimulated region (Fig. 1c), 20 mins before training (see Methods).

Specifically blocking perirhinal cortex input to L1 of S1 severely reduced learning during the first training session (Fig. 1d&e). We quantified learning as the cumulative difference between the number of successful and failed licking responses to μStim (Σ[*hits-misses*]). By this criterion, mice in which the influence of perirhinal axons on L1 of neocortex was suppressed could not associate the water reward with the μStim over the first training session but rather licked in approximately 50% of the trials (average learning score 0.48±0.06 normalized to the total number of trials, n=6 in ctrl versus −0.03±0.08, n=7 in hM4Di/CNO-treated mice; Wilcoxon rank-sum test, p=0.0047). Note, CNO alone^15^, i.e. without expression of hM4Di, had no effect on learning (n = 3, Wilcoxon rank-sum test, p=0.4; Extended Fig. 2). In contrast to control animals, untrained animals rarely responded to μStim (−0.54 ± 0.05, n=5 in untrained mice, Wilcoxon rank-sum test, p=0.0043; Fig. 1d&e). After 3 sessions, trained animals had improved learning scores (0.87±0.04 at session 3, Wilcoxon rank-sum test, p=0.02) and this was not affected by suppression of the perirhinal influence on S1 (0.84±0.06, n=3 in CNO-treated trained mice, Wilcoxon sign-rank test, p=1; Fig. 1e and Extended Fig. 2). This suggests that perirhinal cortex is involved in early memory formation but does not affect perception of the μStim per se. The second order somatosensory thalamic area, POm, also projects to L1 in S1 and has been implicated in different learning paradigms^3–6,16^. To examine the influence of POm input in μStim task, this time we expressed hM4Di receptors in POm in Gpr26-cre transgenic mice^17^. Suppression of this projection from POm slightly affected learning, however the effect was not significant (0.25±0.05, n=7, Wilcoxon rank-sum test, p=0.18; Extended Fig. 2). Taken together, these results show that the influence of the perirhinal cortex on L1 of the neocortex is crucial for learning the μStim detection task.

The results of the chemogenetic experiments (Fig. 1) imply that activity in perirhinal cortex influences activity in S1. Since the μStim electrode was most effective when placed in L5^12^, we reasoned that the stimulation at least affected L5 neurons. Furthermore, L5 neurons have been implicated in perceptual detection tasks and it has been recently shown that the output of these neurons depends partly on the activation of their apical dendrites that project into L1^18^ where the perirhinal inputs arrive. We confirmed ex vivo that perirhinal inputs arriving in L1 synapse on to the tuft dendrites of L5 pyramids (Extended Fig. 3). To investigate the influence of PRh input in S1 activity we made juxtacellular recordings from L5 in S1 in the same region as the μStim (Fig. 2a, left). We recorded activity from S1 during learning with and without chemogenetic suppression of perirhinal input to the cortical L1. As in the purely behavioral experiments (see Fig. 1), hM4Di/CNO-treated animals also did not learn the task over the first session (0.1±0.04, n=4, Wilcoxon rank-sum test against control, p=0.04). Both the baseline (1 s before μStim) and post-stimulus (0.5-2.5 s after μStim) AP firing rates in L5 pyramidal neurons of these animals (n=4, 52 cells, 826 trials) was significantly reduced in comparison to control animals (n=2, 28 cells, 706 trials) treated with CNO only (Wilcoxon rank-sum test, p<0.001 for both baseline and post-stimulus; Fig. 2b-d). These results refer to ‘Hit’ trials where the animals responded correctly to μStim although we found analogous results in ‘Miss’ trials (Extended Fig. 3). There was no significant difference in firing rate in control animals between baseline and post-stimulus activity and a slight but significant reduction in hM4Di/CNO-treated animals (see also Extended Fig. 3).

**Figure 2.**
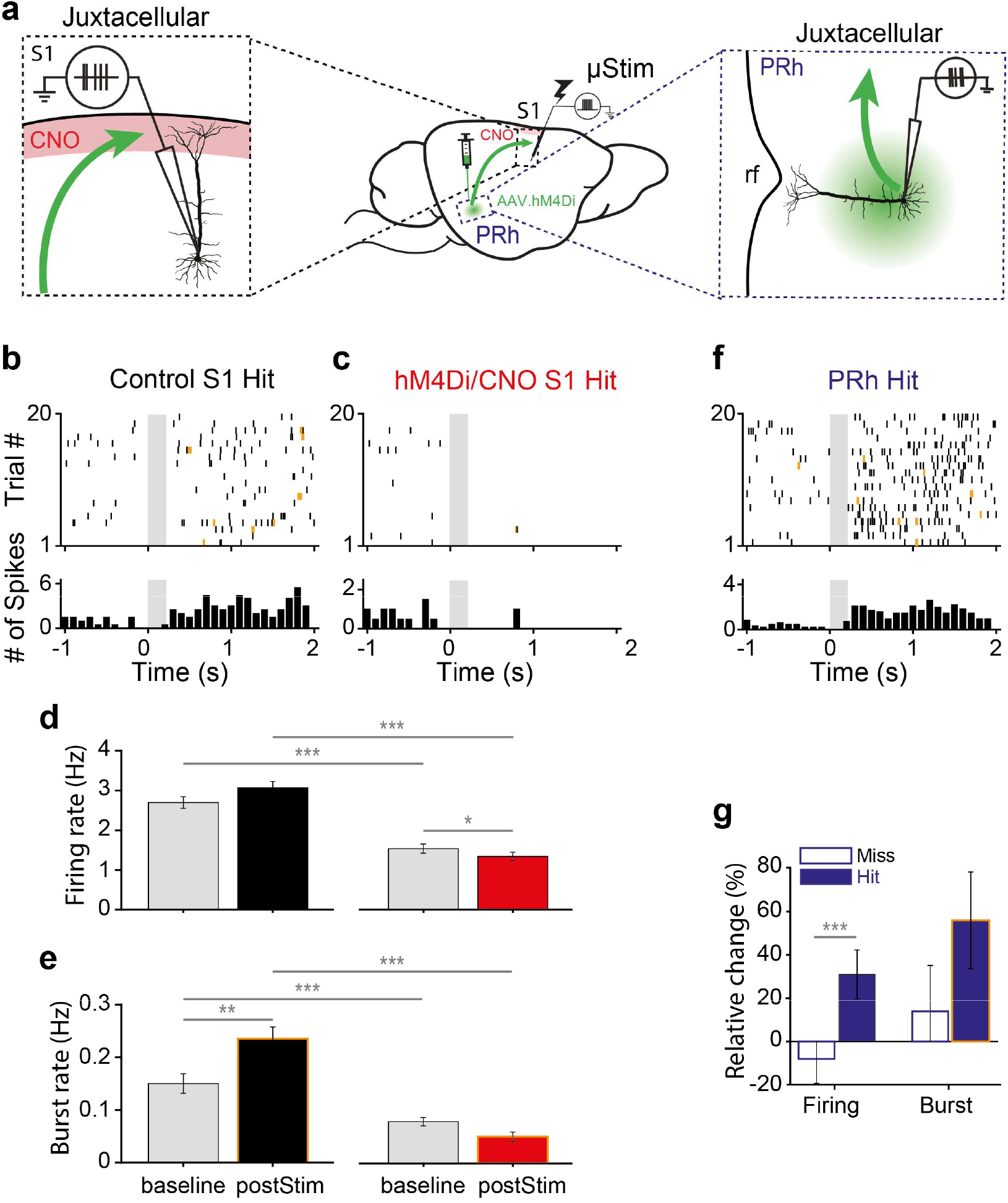
Learning-induced burst firing in layer 5 pyramidal neurons in S1 is PRh-dependent. **a**, Schematic of juxtacellular recording of L5 pyramidal neurons in S1 (left) and deep layer pyramidal neurons in PRh (right). Red shade: CNO targeted area, rf: rhinal fissure. **b, c**, Representative raster plot (upper) and PSTH (lower) during hit trials in L5 pyramidal neurons in control S1 and hM4Di S1, respectively. Bursts are marked by yellow ticks in the raster plot. Gray box: μStim. Note that y-axis scales differ for visibility. **d, e** Firing rate and burst rate during hit trials in control S1 and in hM4Di S1, respectively. **f**, Representative raster plot (upper) and PSTH (lower) during hit trials in perirhinal neuron. **g**, Relative change of firing rate and burst rate during miss and hit trials from PRh neurons. *p<0.05, **p<0.01, ***p<0.001.

Previously, we found that animals are biased to respond to irregular firing patterns in animals trained on a μStim task^19^ and that burst firing correlates with perceptual detection^18^. We therefore examined burst firing of the same cells during learning in control and hM4Di/CNO-treated animals where learning was blocked. Here, we found that blocking of learning via suppression of perirhinal input significantly decreased both baseline and post-stimulus burst rate (Wilcoxon rank-sum test, p<0.001). Interestingly, the burst rate following μStim compared to baseline was greatly increased in control animals (Wilcoxon sign-rank test, p=0.001) but not in hM4Di/CNO-treated animals and only in ‘Hit’ trials (see Extended Fig. 3). We conclude that perirhinal input to L1 mediates learning-related increases in excitability and burst firing in neocortical L5 neurons.

What information does perirhinal cortex convey during μStim learning? To investigate this, we examined the firing of deep layer neurons in perirhinal cortex (Fig. 2a, right). We found that perirhinal neurons responded robustly to hit trials but not to miss trials after μStim in S1 (Fig. 2f&g; n=6 animals, n=287 trials in 28 neurons; firing rate: miss −8±11.2% versus hit 30.9±11.2%, p<0.001, Wilcoxon rank-sum test). Moreover, the responses in perirhinal cortex included an increase in burst rate compared to baseline only during hit trials (Wilcoxon sign-rank test. p<0.001). However, the relative change of burst rate between miss and hit trials was not significant (Fig. 2f&g; burst rate miss 14 ± 21% versus hit 55.9±22.3%, p=0.1, Wilcoxon rank-sum test). This shows that the perirhinal cortex signals information related to ‘Hit’ trials primarily via increased AP firing during learning.

It has recently been shown that memory formation is accompanied by an increase in slow cortical oscillations^20–23^. We therefore also analyzed the local field potential (LFP) signals, taken from the same recordings in S1 and perirhinal cortex to assess cortical oscillations during learning. Theta power (4 - 8 Hz) in S1 was significantly higher in trained versus untrained animals (Extended Fig. 4; Wilcoxon rank-sum test, p<0.0001). Analogously, in perirhinal cortex, we found a significant increase in the theta power in ‘Hit’ compared to ‘Miss’ trials during learning (Extended Fig. 4; Wilcoxon rank-sum test, p=0.002). These results suggest that elevated theta power in perirhinal cortex correlates to a transition to elevated theta in response to μStim in S1 in expert animals.

Depolarization of the apical dendrites in L5 pyramidal neurons is shown to reliably lead burst firing behaviour^24–26^. Since learning correlated with an increase in burst firing in these neurons (Fig. 2e) that was dependent on perirhinal input to L1, we hypothesized that the mechanism of learning-induced bursting might involve an enhancement of synaptic influence to the tuft dendrites. We therefore examined Ca^2+^-dependent activity in the apical dendrites of L5 neurons in S1 using 2-photon microscopy in trained animals. To do this, we expressed GCaMP6f in Rbp4-Cre transgenic mice^27^ and imaged at a depth of ~200 μm, the region of the apical dendrite known for initiation of dendritic Ca^2+^ activity^7^ (Fig. 3a-c). Calcium transients measured from 1 s before the μStim until 3 s after the μStim in 318 dendrites (Fig. 3d; n = 4 mice), revealed three populations with distinct fluorescence profiles (Fig. 3e). A small population (10%, “ON” dendrites) of dendrites showed substantial increases in fluorescence following μStim with another population (37%, “OFF” dendrites) of dendrites showing reduced Ca^2+^ fluorescence. The rest were not responsive to μStim (53%, “NR” dendrites).

**Figure 3.**
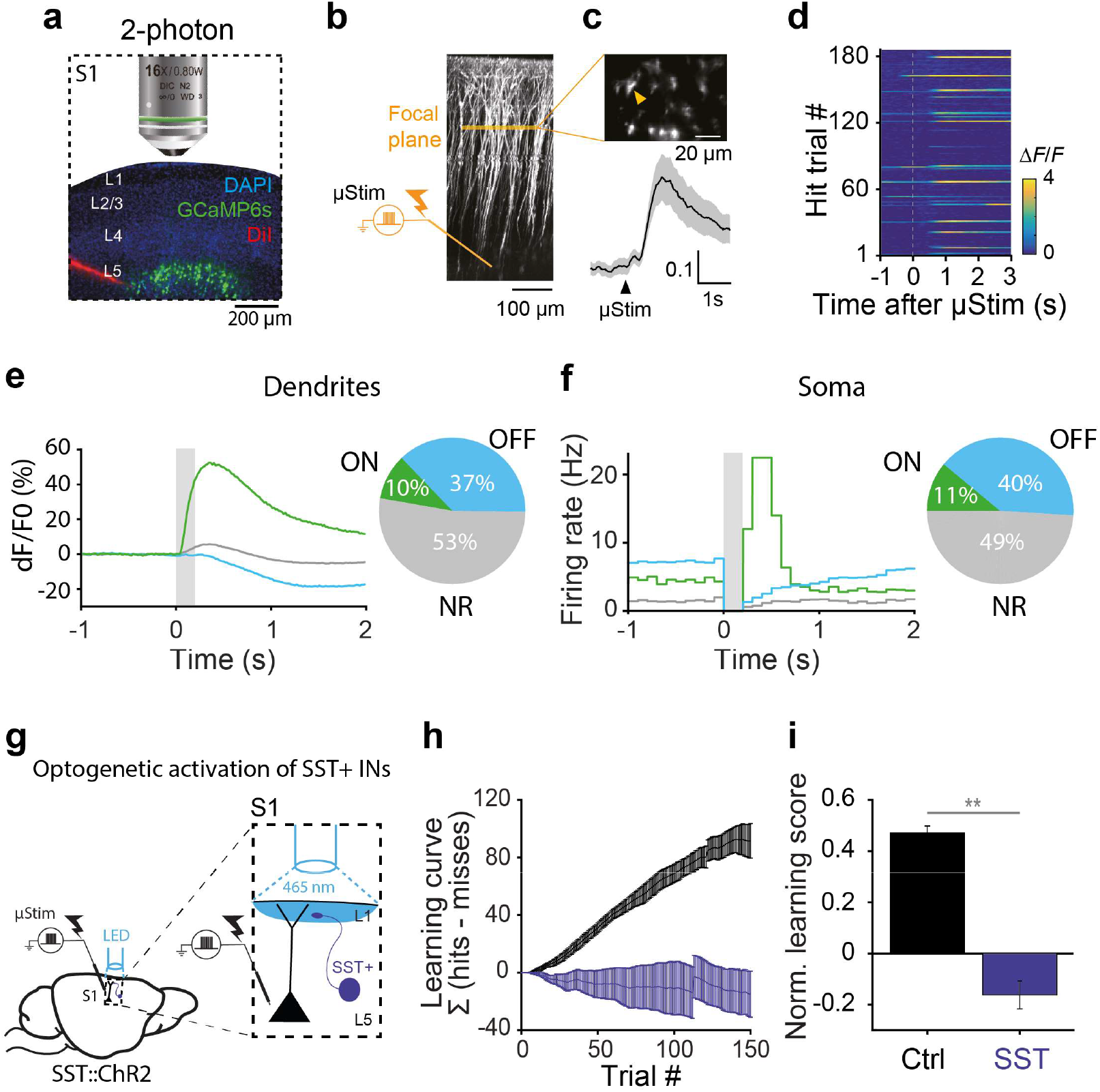
Dendritic activity-dependent emergence of distinct L5 subpopulations after learning. **a**, Two-photon Ca^2+^ imaging from the apical dendrites of L5 pyramidal neurons in Rbp4-cre mice during μStim task. DiI shows the location of μStim electrode. **b**, Z-stack image of recorded dendrites and μStim electrode in L5. **c**, Horizontal imaging plane (upper) (~200 μm from pia) and average Ca^2+^ responses (lower) for all trials from a dendrite marked with a yellow arrow. **d**, Ca^2+^ responses in an apical dendrite marked in c. during 180 trials of μStim task. **e**, Left, Average peri-stimulus time Ca^2+^ responses in ON, OFF and NR dendrites (total n=318 dendrites) during hit trials (See Methods for classification criteria). Gray box: μStim. Right, the fraction of ON, OFF and NS dendrites. **f**, Left, Average PSTH of L5^ON^, L5 ^OFF^ and L5^NR^ neurons (total n=272 cells) during hit trials (See Methods for classification criteria). Gray box: μStim. Right, the fraction of L5^ON^, L5 ^OFF^ and L5^NR^ neurons. **g**, Schematic of optogenetic activation of SST+ interneurons during μStim task in SST::ChR2 mice. Blue light (465 nm) was shed on the surface of the craniotomy (See Methods). Inset, magnified view of the targeted circuit in S1. **h**, Cumulative learning curve of control (n=6, black) and SST::ChR2 mice (n=6, dark blue) during the first session. **i**, Normalized learning score of control and SST::ChR2 mice at the first session. Wilcoxon rank-sum test, ** p<0.01.

We found similarly distinct and stereotypical output firing patterns in L5 neurons using juxtacellular recordings from trained animals (Fig. 3f; see Extended Fig. 5 for examples). In 11% of cells we saw a sudden and marked increase in firing (21.44±42.16 Hz) briefly following μStim (Fig. 3f; L5 “ON” cells). In another population consisting of 40% of neurons, there was a decrease in firing (−6.39±4.93 Hz) immediately following the μStim (Fig. 3f; L5 “OFF” cells). In most of the cells (49%), we observed no response to μStim (L5 “NR” cells). Interestingly, the baseline firing rate in L5 ON and L5 OFF cells was significantly higher than in NR cells (Wilcoxon rank-sum test, NR vs. ON: p < 0.0001, NR vs. OFF: p < 0.0001). In contrast to expert animals, we observed low firing rates over all neurons in untrained animals (Extended Fig. 6). Most L5 neurons in untrained animals did not respond to μStim at all (95%, n=63/66 cells) with a small population (5%, n=3/66 cells) responding with a small increase (6.9±6.30 Hz) briefly after μStim. Taken together with the 2-photon dendritic recordings, we conclude that learning enhances the responsiveness of a small population of L5 pyramidal neurons to apical dendritic input.

To test whether dendritic activity influences learning, we optogenetically activated dendrite-targeting inhibitory neurons during the μStim training. Previous studies have implicated somatostatin (SST) positive interneurons in suppressing plasticity and learning via dendritic inhibition^5,6,8,28,29^. We reasoned that if the same circuitry is activated during the μStim detection task, activating SST neurons with channelrhodopsin2 (ChR2) should also impair learning. We activated SST neurons during training in SST::ChR2 mice using a 500 ms light pulse starting 300 ms before μStim (Fig. 3g). This abolished learning in a manner almost identical to removing the influence of perirhinal input to L1 (SST: −0.16 ± 0.05, Wilcoxon rank-sum test, p=0.002; Fig. 3h&i). Notably, continued activation of SST neurons through subsequent training sessions prevented learning over several sessions, unlike block of perirhinal input in which the animals eventually became experts (Extended Fig. 7). Altogether these results suggest that the emergence of a population of neurons underlying learned behavior in the μStim task depends on a dendritic mechanism.

The correlation between both bursting and dendritic activity with learning suggests that bursting might underlie memory retrieval in cortical neurons. In order to test this hypothesis we devised another learning paradigm in which we first trained animals to respond expertly to μStim and then manipulated the firing of single neurons in S1 using single-cell stimulation (“nanostimulation”^12,19,30^) via a juxtacellular electrode (Fig. 4a). Expert animals were significantly more likely to lick for reward if bursts of APs (80–120 Hz) were elicited in a single L5 pyramidal neuron of S1 compared to false-positive trials where no current was injected. However, response rate to a train of regularly spiking APs (30–50 Hz) was not significantly different from false-positive rate (Fig. 4b&c; Hit rate; false-positive: 25.94±3.6%, regular: 28.11±4.07%, burst: 31.47±4.24%, n=27 cells, one-sided paired t-test, p=0.03). This indicated that burst firing increased the downstream readout of the firing of a single L5 pyramidal neuron leading to successful behavior. Since the learned behavior could be recovered by burst firing in single pyramidal neurons, these data suggest that burst firing observed in L5^ON^ neurons might be extremely effective in memory recall.

**Figure 4.**
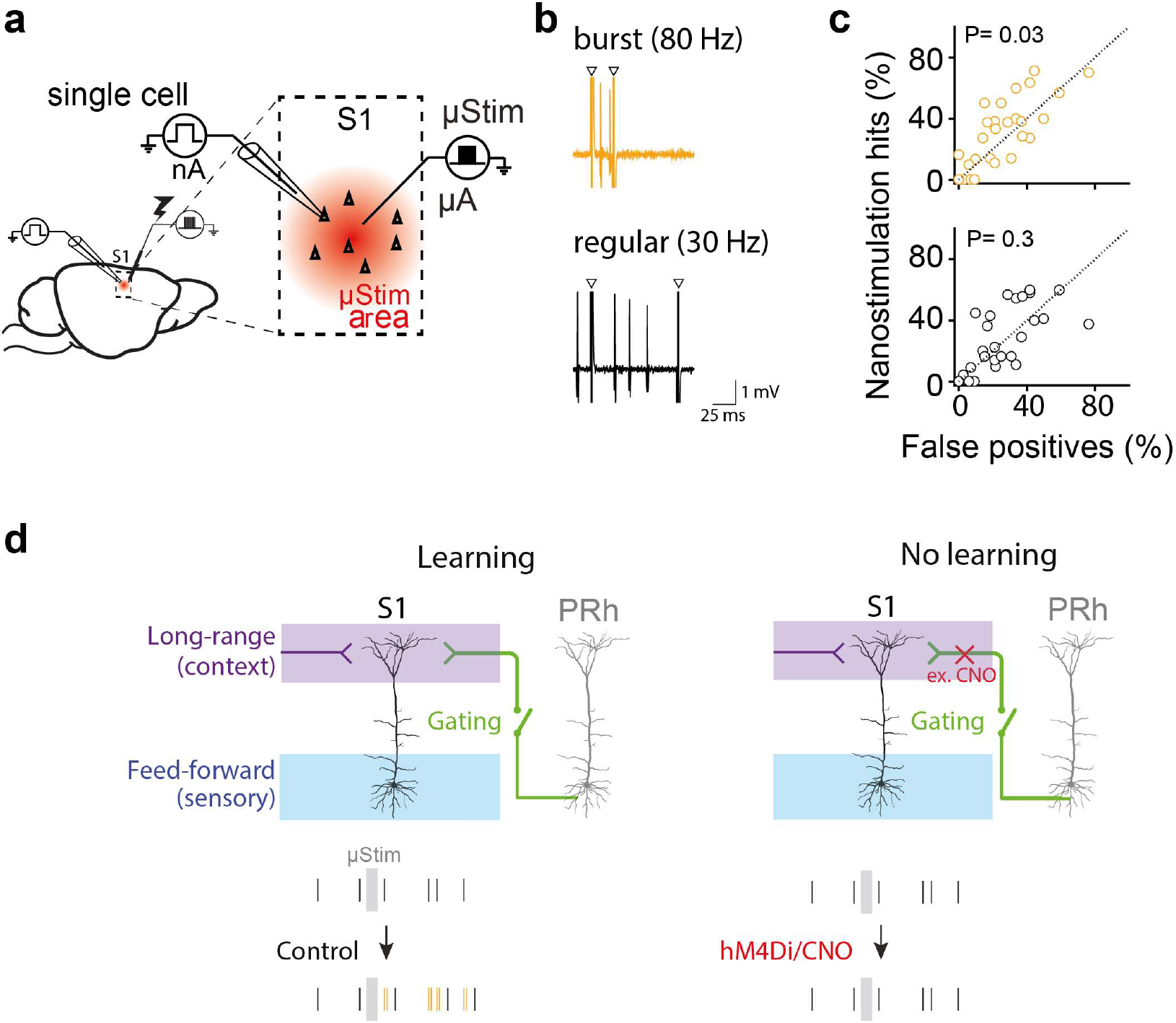
Burst firing in single L5 pyramidal neurons can retrieve learned behavior. **a**, Schematic of single-cell stimulation. Inset, μStim stimulates a population of neurons surrounding electrode (red shade) and glass electrode stimulates a single neuron (triangle). **b**, Current injection protocol to induce either regular AP firing (30 Hz, black) or high frequency (80-120 Hz, yellow) bursts in single cells. **c**, Response rates (hits) for regular AP firing (black) or burst firing (yellow) trials versus false-positive trials (n=27 cells). One-sided paired t-test. **d**, Gating theory of memory formation in cortex. PRh inputs to L1 gate long-range inputs that modulate the firing mode of L5 pyramidal neurons in S1 and learning.

Overall, we have shown that the perirhinal connection to L1 of neocortex is crucial for learning a μStim detection task and involves the conversion of neurons in neocortex to high-firing, burst mode correlated with an increase in dendritic activity (Fig. 4d). This implies that the apical dendrites of L5 neurons are the locus of plasticity related to memory consolidation. This idea is corroborated by our previous study where we showed that stimulus detection in S1 was dependent on dendritic activity in trained animals^18^ and was disrupted by inhibiting this activity. In addition, dendritic activity is shown to be generated by feedback signals from other cortical areas^7,9^ and is enhanced during learning^31^, suggesting that perirhinal input to L1 might serve as a gating signal for the enhancement of cortico-cortical feedback inputs (Fig. 4d). We conclude that the medial temporal input to the neocortex controls learning through a process in L1 that is encoded by dendritic calcium promoting burst firing as the neural signature of memory recall.

## Methods

### Animals

All experiments and procedures were approved and conducted in accordance with the guidelines given by Landesamt für Gesundheit und Soziales Berlin. The following animal lines were used in this study: C57BL/6J wild-type mice, Gpr26-cre transgenic mice^17^, SST::ChR2 transgenic mice (SST-IRES-Cre mice (JAX stock #018973) were crossed with Ai32 mice (JAX stock #024109))^32^ Rbp4-cre transgenic mice^27^ and Wistar rats (Charles River). Male animals were used except for 2 Rbp4-cre mice. The animals were housed in reversed 12 h light/dark cycle (light on between 21:00 and 09:00) and all the behavioral experiments were performed during dark period of the cycle.

### Retrograde and anterograde tracing

For the retrograde labeling of S1 projecting perirhinal neurons, Fast Blue (25% in dH_2_O, Polysciences) soaked in a sterile piece of tissue was applied onto the surface of S1 for 10 min. Incubation time was 7 days before transcardial perfusion. For anterograde tracing and optogenetic *ex-vivo* experiments AAV-hSyn-hChR2(H134R)-EYFP (Penn Vector Core) was injected in the PRh of > 2 weeks old C57/BL6 mice. Anesthesia was induced and maintained with isoflurane at 5% and 2%, respectively. Mice were placed in a stereotaxic frame and craniotomies were performed using stereotaxic coordinates: anterior-posterior axis (AP) −1.8 mm, medial-lateral axis (ML) ± 4.1 mm, DV −4.2 mm from bregma. Injections were carried out using graduated pipettes broken back to a tip diameter of 10-15 μm, at a rate of ~ 0.025 μl/min for a total volume of 0.05-0.07 μl. Incubation time was at least 3 weeks before transcradial perfusion or *ex-vivo* experiment.

### YFP fluorescence analysis

AAV-hSyn-hChR2(H134R)-EYFP containing acute brain sections were imaged using an Olympus BX51 Microscope with a 4x objective. Fluorescence intensity was quantified with ImageJ software by plotting a line profile across the cortical layers that calculates the brightness value. The average gray value of all images was then normalized to the negative SEM of the lowest grey value across the average line profile.

### *Ex-vivo* electrophysiology

After 3-4 weeks of virus expression, sagittal or coronal slices (300 μm thick) were prepared from 35-50 day old C57/BL6 mice. Whole-cell patch-clamp recordings were performed from visually identified layer-5 pyramidal neurons using infrared (IR) Dodt-gradient contrast video microscopy. The extracellular solution contained 125 mM NaCl, 25 mM NaHCO3, 25 mM Glucose, 3 mM KCl, 1.25 mM NaH2PO4, 2 mM CaCl2, 1 mM MgCl2, pH 7.4 at ~33 ° C. The intracellular solution contained 115 mM K+-gluconate, 20 mM KCl, 2 mM Mg-ATP, 2 mM Na2-ATP, 10 mM Na2-phosphocreatine, 0.3 mM GTP, 10 mM HEPES, 0.05 mM Alexa 594 and biocytin (0.2%), pH 7.2. Whole-cell voltage recordings were performed from the soma (4-6 MΩ) using a Multiclamp 700b (Molecular devices) amplifier. Data was acquired with an ITC-18 board and analyzed using Igor software. Optogenetic synaptic stimulation was performed via an LED (470 nm) (2 ms pulses) located in L1 around the tuft dendrite. To activate hM4D_i_ receptor, Clozapine-N-Oxide (CNO) (Tocris Bioscience) was bath applied (final concentration 10 μM).

### Chemogenetic manipulation of PRh axonal activity

Mice (> 4 weeks) were anesthetized with ketamine/xylazine (13 mg kg^−1^/1mg kg^−1^) by intraperitoneal injection (i.p.). Animals were kept on a thermal blanket during entire surgery and recovery. Lidocaine (1%, wt/vol, AlleMan Pharma) was injected around the surgical site before the scalp incision. The periosteum was removed and small craniotomy was made on the injection sites. Injection coordinates for PRh were AP −1.8 mm, ML ± 4.2 mm, DV −4.2 mm and for POm were AP −2 mm, ML +1.2 mm, DV −3.0 mm from bregma. AAV1/2-hSyn1-hM4D(Gi)-mCherry-WPRE-hGHp(A) (Viral Vector Facility of the University of Zurich) was injected either bilaterally (n=16 for PRh and n=7 for Pom) or unilaterally (n=3 for PRh) (0.15 –0.20 μl per side). Further experiments were performed after 3 weeks of expression.

In order to activate hM4Di receptor, CNO dissolved in extracellular solution (10 μM) was applied into superficial layers of S1 (initial depth at 150 μm and over the day injection pipette was advanced (up to 300 μm) in order to compensate tissue growth on the craniotomy) at least 20 min before the microstimulation training. CNO was applied into two adjacent sites (150 μl each) of the craniotomy to maximize the CNO diffusion area.

### Headpost implant and head-restraint habituation

A lightweight aluminum head-post (mouse) or a metal bolt (rat) was implanted on the skull of the animal under ketamine/xylazine anaesthesia (13 mg kg^−1^/1mg kg^−1^ for mice, 100 mg kg^−1^/5 mg kg^−1^ for rats, i.p.). For mice used in chemogenetic experiments, the implantation was performed > 10 days after viral injection. After the scalp and periosteum were removed, a thin layer of light curing adhesives (OptiBond, Kerr and Charisma, Kulzer) was applied to the skull. A head-post was fixed on the skull on the left hemisphere with a dental cement (Paladur, Heraeus Kulzer).

Head-restraint habituation began > 3 days after the head-post implantation. Habituation time at the first day was 5 min and then gradually increased each day until the animal sat calmly for 1 h. Animals were water restricted from the second day (1 ml/day) of the habituation and then trained to receive the saccharin (Sigma-Aldrich) water (0.5% for mice and 0.1% for rats) from the licking port. Licking was monitored using a piezo-based sensor attached to the licking port. Weight and health of the animal were monitored daily. Habituation for head restraint and licking typically took 5 days.

Two to three days before the microstimulation training or/and juxtacellular recording, 1.5 mm x 1.5 mm craniotomy was made on the right barrel cortex centered at AP 1.25 mm and ML 3.75 mm from bregma for mice and AP 2.5 mm and ML 5.5 mm from bregma for rats. For perirhinal cortex recording in rats, craniotomy was made on AP 4.5 mm and ML 5.0 mm from bregma. Then a recording chamber was implanted for chronic access to this region. The dura was left intact and the craniotomy was covered with silicon (Kwik-Cast, World Precision Instruments).

### Optogenetic manipulations

For optogenetic activation of SST neurons, SST::ChR2 transgenic mice that express ChR2 in SST-positive cells were used. Photostimulation light (465 nm, 2 mW, 500-ms pulse starting at 300 ms before stimulus onset) was delivered via the optic fiber placed above the craniotomy. To prevent the mice from distinguishing photostimulation trials from control trials using visual cues, the recording chamber was covered with a black rubber to prevent light leakage from photostimulation into the animals’ eyes.

### Microstimulation detection task

Animals were trained to perform microstimulation task as described elsewhere^12,19,33^. Briefly, animals were trained to respond with tongue lick to a 200 ms train of microstimulation pulses applied to barrel cortex (40 cathodal pulses at 200Hz, 0.3 ms pulse duration) through a tungsten microelectrode (Microprobes) in depth of ~700 μm (mice) or ~1500 μm (rats) from pia and presented at random intervals. In the first session, initial intensity of 160 μA pulses were injected into the cortex and paired with a drop of water reward (pairing period). After 5 pairings, testing began where animals were rewarded only if they lick the licking port within 100 to 1,200 ms after stimulus onset. Tongue lick responses were detected with piezo-based sensor (mice) or beam breaker (rats). The time of the first lick after stimulus onset was taken as the reaction time. To encourage animals to use a nonconservative response criterion, we only mildly punished licks in the interstimulus interval with an additional 1.5 s delay to the next stimulus presentation. Once animals reached 80% hit rate, pulse intensity was gradually decreased during and over the sessions until it reached 10 μA (mice) or 5 μA (rats). Control mice reached to 10 μA within 3-5 days of training. Expert in this study means animals who performed the task with >80% hit rate at the 10 μA (mice) or 5 μA (rats).

### *In vivo* juxtacellular recording

Following head-restraint habituation, juxtacellular recordings were performed from deep layer neurons from S1 and PRh in awake head-fixed animals during μStim detection task. The glass pipette (4–8 MΩ) for juxtacellular recording during microstimulation task was filled with extracellular solution containing: 135 mM NaCl, 5.4 mM KCl, 1.0 mM MgCl2, 1.8 mM CaCl2 and 5 mM HEPES (pH 7.2). The juxtacellular signal was amplified and low-pass filtered at 3 kHz by a patch-clamp amplifier (NPI) and sampled at 25 kHz by a Power1401 data acquisition interface under the control of Spike2 software (CED). For PRh recording in rats, the pipette was inserted with 17° toward lateral and 50° toward anterior. The mean depth in juxtacellular recording of S1 in mice was 1156.8±25.56 mm and of PRh in rats was 6339.64±122.07 mm, which is likely an overestimate of the true depth due to oblique penetrations and dimpling.

### Two-photon Ca^2+^ imaging

For *in vivo* two-photon calcium imaging, AAV2/1-Syn-Flex-GCaMP6f-WPRE (Penn Vector Core) was injected through a glass pipette (tip diameter, 5–10 μm) into the left S1 barrel cortex on the basis of stereotaxic coordinates (AP −1.5 mm and ML 3.2 mm from bregma). A single injection (100 nl) was made at 700 μm deep from the pial surface. Three weeks after the injection, a 3-mm craniotomy was made over the injection site and sealed with a 3-mm glass coverslip (#1) with cyanoacrylate glue. A light-weight head-post was fixed on the skull in the right hemisphere with light-curing adhesives and a dental cement. Habituation of mice to head restraint and following imaging experiment begin 4 weeks after the virus injection.

Imaging from behaving mice was performed with a resonant-scanning two-photon microscope (Thorlabs) equipped with GaAsP photomultiplier tubes (Hamamatsu Photonics). GCaMP6f was excited at 940 nm (typically 30–40 mW at the sample) with a Ti:Sapphire laser (Mai Tai eHP Deep See, Spectra-Physics) and imaged through a 16×, 0.8 NA water immersion objective (Nikon). Full-frame images (256 × 256 pixels) were acquired from apical dendrites of L5 neurons expressing GCaMP6s at a depth of 150–200 μm at 58.6 Hz using ScanImage 4.1 software (Vidrio Technologies). Tungsten electrodes for microstimulation was inserted through the access port on the chronic glass window.

### Histology

Animals were perfused transcardially with phosphate-buffered saline (PBS) followed by 4% paraformaldehyde (PFA). Brains were removed and post-fixed for > 24 h. Coronal sections (150 μm thick) were collected using a vibratome. For DAPI staining, NucBlue (Invitrogen) was applied to sections in PBS for 10 min. The sections were imaged with an epifluorescence microscope or a confocal microscope.

### Single-Neuron Stimulation Detection Task

Once rats performed at current intensities below 5 μA on 2 consecutive days, we switched to single-cell stimulation experiments, as previously described^12,33^. Briefly, the animals were head fixed during the task, and waited for the microstimulation/nanostimulation detection task to begin, which it did when a neuron was found. The glass pipette for juxtacellular single-cell stimulation and recording was glued to a tungsten microelectrode used for microstimulation at a distance of ~70 mm, as described elsewhere^12,33^. The glass pipette was filled with intracellular solution containing: 135 mM K-gluconate; 10 mM HEPES; 10 mM Na2-phosphocreatine; 4 mM KCl; 4 mM MgATP; and 0.3 mM Na3GTP (pH 7.2). Recording depth was 1902±60.73 mm, which is likely an overestimate of the true depth due to oblique penetrations and dimpling.

During single-cell stimulation trials, a fixed duration square-wave current pulse was injected into a neuron through a glass pipette. Every stimulation sequence contained each step exactly once, while their order was varied pseudo-randomly from trial to trial. To induce a regular spike pattern, we used a single 100 ms DC current step. To elicit burst like spike pattern, brief stimulation duration of 25 ms was used, followed by 1175 ms inhibition at current intensities of 50% used in the nanostimulation, to prevent any further spikes during the stimulation trial. Single-cell stimulation trials, catch trials without current injection and microstimulation trials were pseudo randomly interleaved in series of 6 trials including 3 microstimulation trials, 2 single-cell stimulation trials (each of different duration) and 1 catch trial. All trials were presented at random intervals (Poisson process, mean 3 s). Microstimulation currents were adjusted (range 3-8 μA, mean 4.2 ± 1.1 μA (s.d.)) such that animals performed close to the detection threshold, resulting in an average microstimulation hit rate of 90%.

### Data analysis and statistics

Recorded neurons were separated into putative fast-spiking (FS) interneurons and regular-spiking (RS) pyramidal neurons based on spike half-width and firing rate. Cells with spike half-width lower than 0.5 ms and firing rate higher than 8 Hz were classified to FS. Only RS were used for further analysis.

All the cells and trials recorded over days were pooled together for comparing activity (firing rate or burst rate) changes during miss and hit trials. Bursts were identified as at least two spikes with an inter-spike interval of ≤ 15 ms. Time window between 1 and 0 s before the stimulus ([−1 0] s) was used to calculate the baseline activity and 0.5 – 2.5 s ([+0.5 +2.5] s) after stimulus was used to calculate post-stimulus activity. For firing rate and burst rate change analysis, difference between pre-stimulus frequency and post-stimulus frequency was divided by average pre-stimulus frequency.

All analysis for Ca^2+^ imaging was performed using imageJ and custom written codes in Matlab. Horizontal and vertical drifts of imaging frames due to animal motion were corrected by registering each frame to a reference image based on whole-frame cross-correlation. The reference image was generated by averaging any given consecutive 100 frames in which motion drifts were minimal. Regions of interest (ROIs) for apical dendrites of L5 neurons were manually selected with the help of average intensity and standard deviation projections across movie frames. For each ROI, pixel values inside the ROI were averaged to obtain the time series of Ca^2+^ fluorescence. The extracted signals were corrected for neuropil contamination by subtracting the local, peri-dendritic neuropil signals. Fluorescence change (Δ*F*/*F*_0_) was calculated as (*F* − *F*_0_)/*F*_0_, where *F*_0_ was the baseline fluorescence value in the ROI throughout the whole imaging session.

For local field potential (LFP) analysis, juxtacellularly recorded voltages were band-pass filtered at 4-30 Hz and a power spectrum was calculated using the Stockwell Transform^34,35^, over a 2 s period before stimulus onset and 5 s period afterwards. In order to avoid potential artifacts caused by stimulation, the analysis was restricted to microstimulation or nanostimulation trials where the following trial occurred more than 5 s after stimulus onset. The power spectrum was calculated separately for individual trials and its absolute magnitude averaged within the different response categories (hits and misses). To obtain the population spectra for the different response categories the power spectra of individual trials were then averaged.

For the classification of cells, peri-stimulus time-histograms (PSTHs) were calculated for each cell by averaging spikes in time bins of 100 ms for times within 2 seconds of hit-trials. For each cell, the stationary rate and standard deviation were computed based on the PSTHs in the period [−2,0] s. Cells were classified to ON cell or OFF cell if PSTHs in the period [0.3,0.4] s was either more than 3*standard deviation (SD) above the stationary rate, or less than 3*SD below, respectively. Other cells were classified as NR cells. Similarly, dendrites were classified into ON, OFF and NR dendrites.

Unless otherwise stated, all values are indicated as mean ± SEM. Shapiro-Wilk test was performed to test normality of the data. For non-parametric test, significance was determined using Wilcoxon signed-rank test within group and Wilcoxon rank-sum test between groups at a significance level of 0.05. No statistical tests were run to predetermine sample size, and blinding and randomization were not performed.

