## Extended data for "Perirhinal input to neocortical layer 1 controls learning"

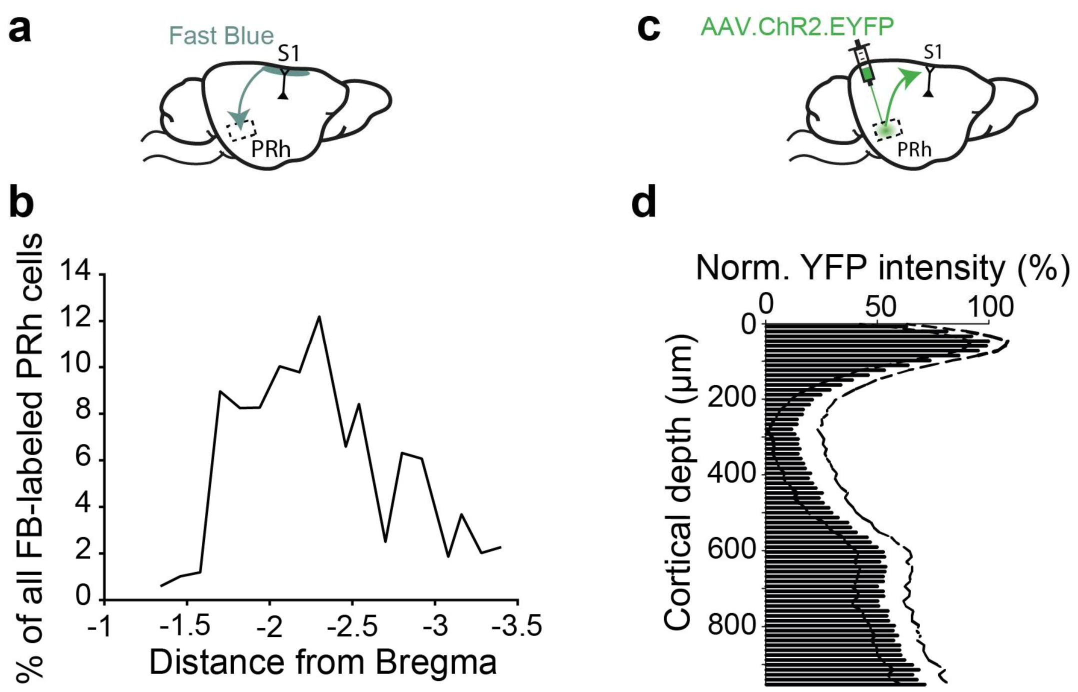


**Extended Data Figure 1 | Quantification of retrograde and anterograde tracing.** **a,** Schematic of retrograde tracing. Fast Blue was applied on the L1 of S1. **b,** Quantification of Fast Blue labelled cells in PRh across anterior-posterior axis. **c,** Schematic of anterograde tracing. AAV.ChR2.EYFP was injected into PRh. **d**, Quantification of YFP intensity across cortical column. YFP intensity was normalized to the lowest mean minus its negative SEM (n=20 sections). Bars (bin size 12.5 µm) and dotted lines represent mean and SEM, respectively.

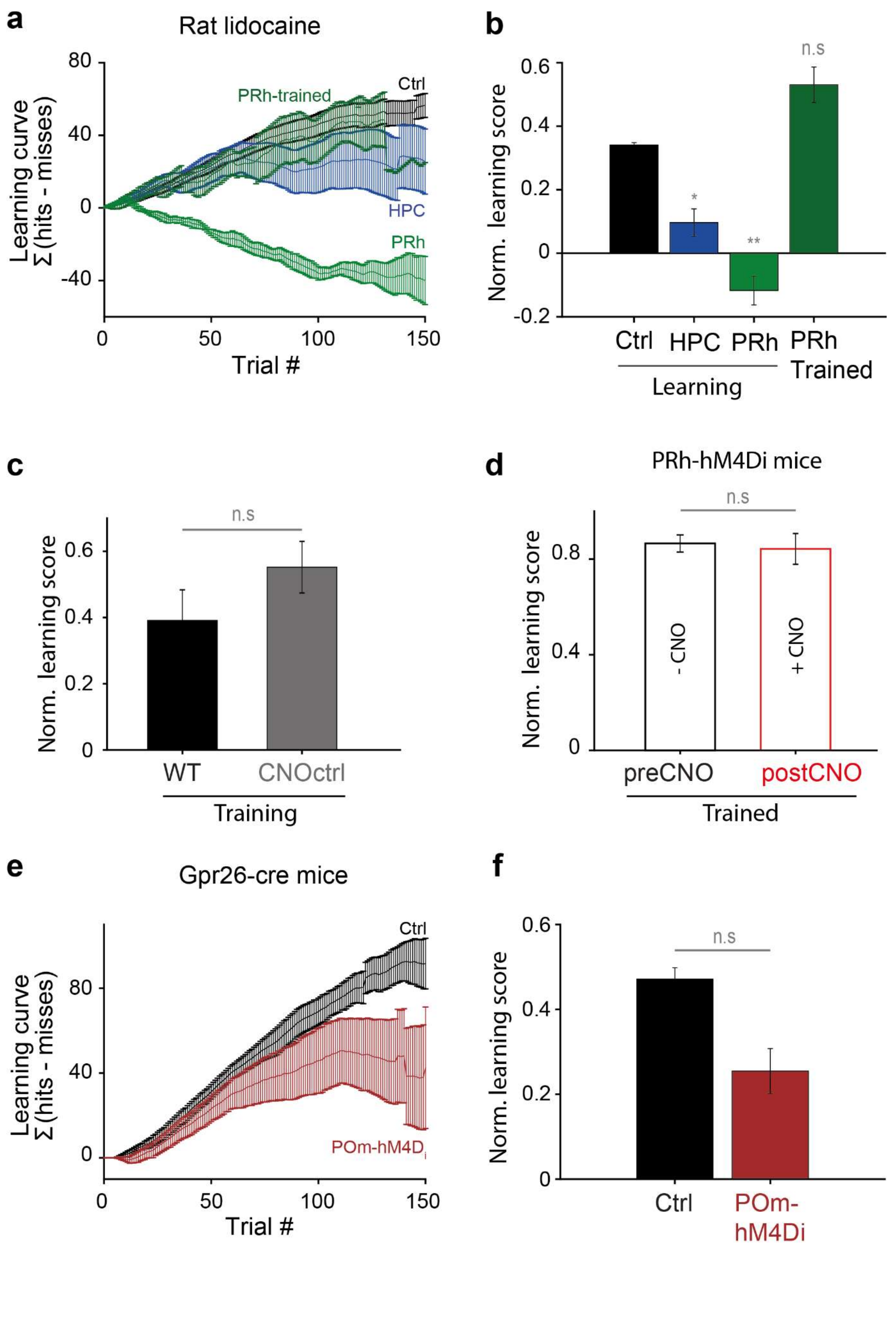
a
Rat lidocaine
Learning curve
Σ (hits - misses)
Trial #
PRh-trained
Ctrl
HPC
PRh
80
40
0
-40
0
50
100
150
b
Norm. learning score
n.s
*
**
0.6
0.4
0.2
0
-0.2
Ctrl
HPC
PRh
PRh
Trained
Learning
c
Norm. learning score
n.s
0.6
0.4
0.2
0
WT
CNOctrl
Training
d
PRh-hM4Di mice
Norm. learning score
n.s
0.8
0.4
0
- CNO
+ CNO
preCNO
postCNO
Trained
e
Gpr26-cre mice
Learning curve
Σ (hits - misses)
Trial #
Ctrl
POm-hM4Di
80
40
0
0
50
100
150
f
Norm. learning score
n.s
0.6
0.4
0.2
0
Ctrl
POm-
hM4Di

**Extended Data Figure 2 | Learning curve of μStim detection task in rodents.**

**a**, Learning was quantified as the cumulative difference between the number of successful and failed licking responses to μStim (Σ[*hits-misses*]). Cumulative learning curve of control rats (black), rats with lidocaine injected to the hippocampus (HPC, dark blue), rats with lidocaine injected to PRh (green) during the first session and rats with lidocaine injected to PRh after learning (dark green). **b**, Normalized learning score (Ctrl, n=31; HPC, n=7; PRh, n=4; PRh expert, n=6). Kruskal-Wallis test, p=0.0052; post-hoc Wilcoxon rank-sum test against Ctrl, HPC: p=0.0460; PRh: p=0.007; PRh expert: p=0.26. **c,** Normalized learning score between wild-type (n=3) and CNO control mice (n=3). Wilcoxon rank-sum test, p=0.4. **h,** Normalized learning score before and after CNO application in trained mice expressing hM4Di in PRh (n=3). Wilcoxon rank-sum test, p=1. **e**, Cumulative learning curve in first session for control mice (black, n=6) and mice with POm axonal suppresion (POm-hM4D$_i$, dark red, n=7). **d**, Normalized learning score. Wilcoxon rank-sum test, p=0.18.

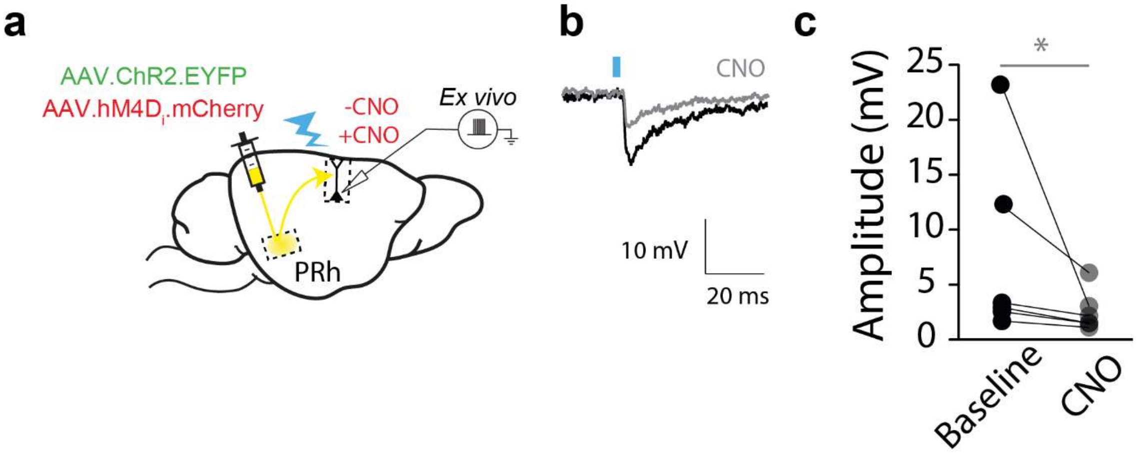


**Extended Data Figure 3 | Cortical pyramidal neurons receive input from PRh.**

**a**, Schematic of *ex vivo* experiments describing virus injection site and optogenetic stimulation of PRh axons in S1. AAV.ChR2.EYFP (green) and AAV. hM4Di.mCherry (red) were co-injected into PRh. **b**, EPSP from a representative cell before and after CNO application in bath. Postsynaptic EPSP induced by light activation (blue line) of PRh axons was reduced after CNO application. **c**, Summary of **f**. Each dot corresponds to a cell (n=6 cells). Baseline 7.64±3.47 mV versus CNO 2.58±0.76 mV, Wilcoxon sign-rank test, p = 0.0313.

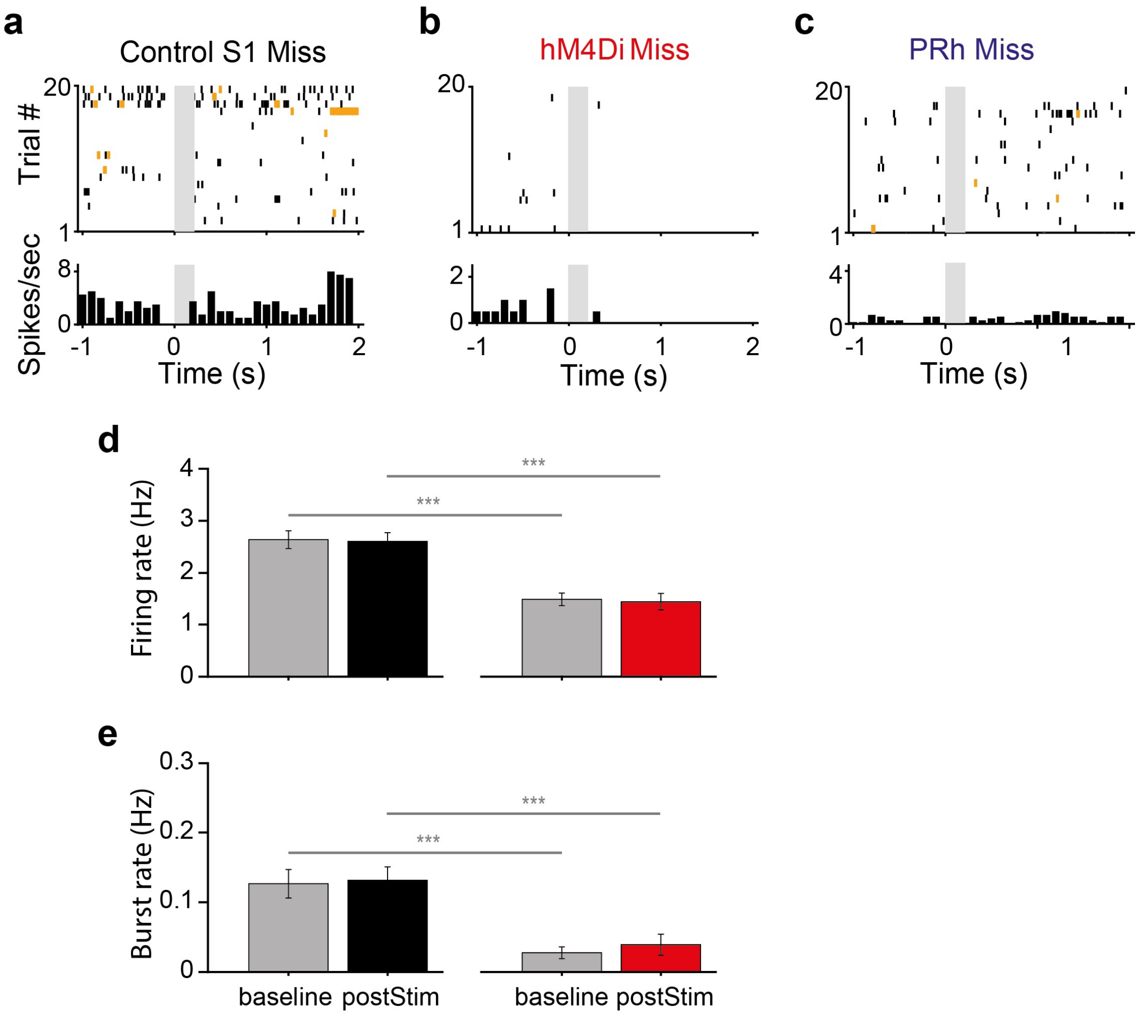


**Extended Data Figure 4 | Somatic activity of S1 in control and hM4Di mice and of rat PRh during miss trials.**

**a, b, c,** Raster plot and PSTH of a representative cell in control S1, hM4Di S1 and control PRh shown in Fig.2 during miss trials. Bursts are marked by yellow ticks in the raster plot. Gray box: μStim. Note that y-axis scales are different for visibility. **d, e,** Firing rate and burst rate during miss trials in control S1 and hM4Di S1, respectively. Wilcoxon rank-sum test, ***p<0.001.

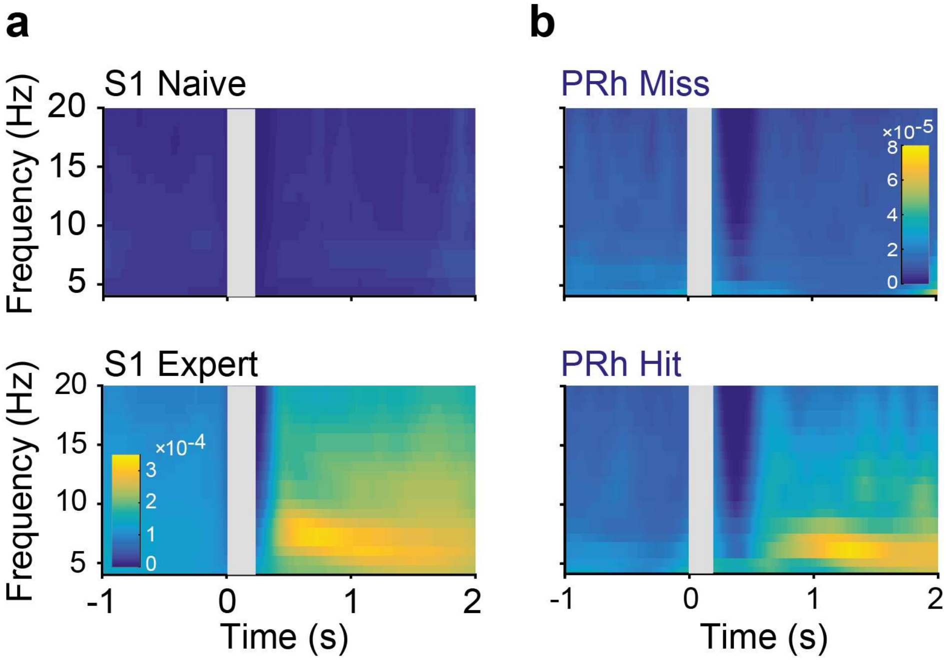


**Extended Data Figure 5 | Local field potential changes in S1 and PRh during μStim task.**

**a,** Time-frequency spectra of local field potential measured in S1 of Untrained and Trained animals. Note the power increase in theta frequency range (4-8 Hz) in expert after μStim (0.2 ± 0.02 arbitrary units (AU) versus 0.03 ± 0.003 AU, $p < 0.0001$). **b,** Time-frequency spectra of LFP measured in PRh during miss and hit trials. Note the power increase in theta frequency range only during hit trials (Hit trials, 0.04 ± 0.01 AU; Miss trials, 0.02 ± 0.002 AU, Wilcoxon rank-sum test, p=0.002).

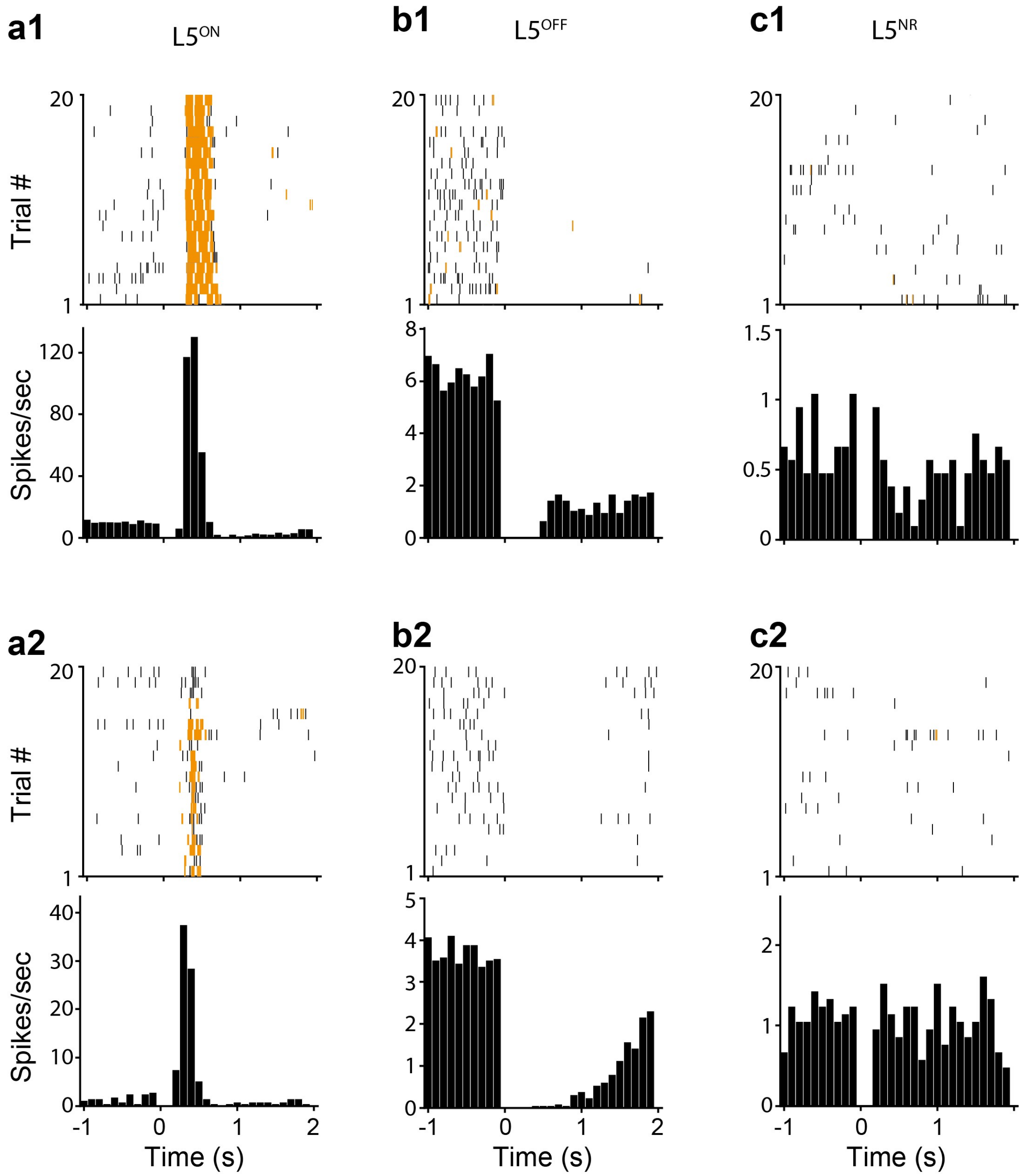


**Extended Data Figure 6 | Raster plot and PSTH of OFF and NR cells.**

**a, b, c** Raster plot (upper) and PSTH (lower) of a representative $L5^{ON}$, $L5^{OFF}$ and $L5^{NR}$ neurons, respectively. Bursts are marked by yellow ticks in the raster plot. Gray box: μStim. Note that y-axis scales differ for visibility.

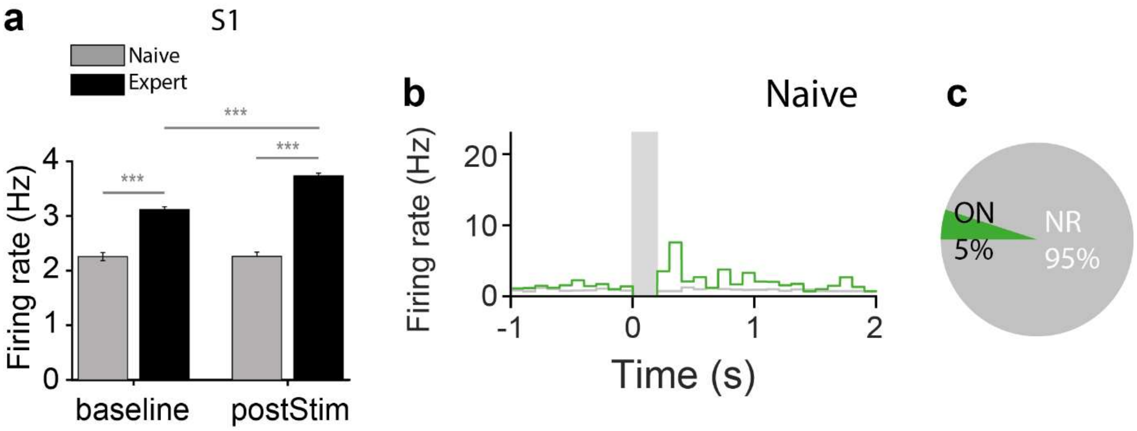


**Extended Data Figure 7 | Increase of somatic activity after learning.**

**a**, Firing rate of S1 L5 pyramidal neurons in Untrained (pre: 2.3±0.07, post: 2.3±0.08, n=70 cells, 2,829 trials) and Trained animals (pre: 3.1±0.04, post: 3.7±0.05, n=278 cells, 20,673 trials) during μStim task. *** p-value < 0.001. **b,** Average PSTH of $L5^{ON}$, $L5^{OFF}$ and $L5^{NR}$ neurons in Untrained animals. **c,** The fraction of $L5^{ON}$, $L5^{OFF}$ and $L5^{NR}$ neurons (total n=66 neurons).

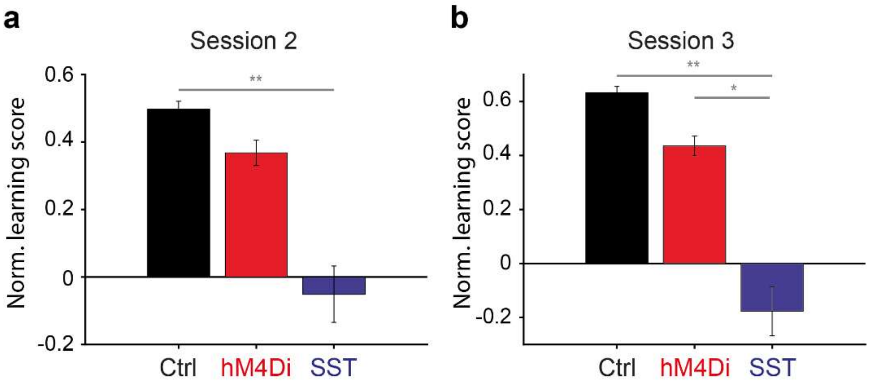


**Extended Data Figure 8 | Persistent impairment of learning in SST::ChR2 mice.**

**a,** Normalized learning score of control (0.5±0.02), hM4Di (0.37±0.04) and SST::ChR2 mice (-0.05±0.08) at second session. Kruskal-Wallis test, p=0.03; post-hoc Wilcoxon rank-sum test, Ctrl vs. hM4Di: p=0.45, Ctrl vs. SST::ChR2: **p=0.009, hM4Di vs. SST::ChR2: p=0.07. **b,** Normalized learning score of control (0.63±0.024), hM4Di (0.44±0.04) and SST::ChR2 (-0.18±0.09) mice at third session. Kruskal-Wallis test, p=0.011; post-hoc Wilcoxon rank-sum test, Ctrl vs. hM4Di: p=0.23, Ctrl vs. SST::ChR2: **p=0.004, hM4Di vs. SST::ChR2: *p=0.04.
